# Intragenic recruitment of NF-κB drives alternative splicing modifications upon activation by the viral oncogene TAX of HTLV-1

**DOI:** 10.1101/676353

**Authors:** Lamya Ben Ameur, Morgan Thenoz, Guillaume Giraud, Emmanuel Combe, Jean-Baptiste Claude, Sebastien Lemaire, Nicolas Fontrodona, Hélène Polveche, Marine Bastien, Antoine Gessain, Eric Wattel, Cyril F. Bourgeois, Didier Auboeuf, Franck Mortreux

## Abstract

The chronic NF-κB activation in inflammation and cancer has long been linked to persistent activation of NF-κB responsive gene promoters. However, NF-κB factors such as RELA also massively bind to gene bodies. Here, we demonstrate that the recruitment of RELA to intragenic regions regulates alternative splicing upon activation of NF-κB by the viral oncogene TAX of HTLV-1. Integrative analysis of RNA splicing and chromatin occupancy, combined with chromatin tethering assays, demonstrate that DNA-bound RELA interacts with and recruits the splicing regulator DDX17 in a NF-kB activation-dependent manner, leading to alternative splicing of target exons thanks to DDX17 RNA helicase activity. This NF-kB/DDX17 axis accounts for a major part of the TAX-induced alternative splicing landscape that mainly affects genes involved in oncogenic pathways. Collectively, our results demonstrate a physical and direct involvement of NF-κB in alternative splicing regulation, which significantly revisits our knowledge of HTLV-1 pathogenesis and other NF-κB-related diseases.

## Introduction

The Human T-cell leukemia virus (HTLV-1) is the etiologic agent of Adult T-cell Leukemia/Lymphoma (ATLL) ^1^, an aggressive CD4+ T-cell malignancy, and of various inflammatory diseases including the HTLV-1-associated myelopathy/tropical spastic paraparesis (HAM/TSP) ^2^. It has long been established that changes in gene expression level participate to the persistent clonal expansion of HTLV-infected CD4+ and CD8+ T-cells, leading ultimately to HTLV-1 associated diseases ^3^. We recently reported that alternative splicing events help to discriminate between ATLL cells, untransformed infected cells and their uninfected counterparts derived from patients ^4^. Alternative splicing of pre-messenger RNAs is a cotranscriptional processing step that controls both the transcriptome and proteome diversity and governs in turn cell fate. Its regulation relies on a complex and still incompletely understood interplay between splicing factors, chromatin regulators and transcription factors ^5,6^. In this setting, the molecular mechanisms of HTLV-1-induced splicing modifications and whether these effects rely on an interplay between transcription and splicing is not known.

TAX is an HTLV-1-encoded protein that regulates viral and cellular gene transcription. TAX also alters host signaling pathways that sustain cell proliferation and lead ultimately to cell immortalization ^7^. The Nuclear factors κB (NF-κB) signaling pathway is the most critical target of TAX for cell transformation ^8^. The NF-κB transcription factors (RELA, p50, c-Rel, RelB, and p52) govern immune functions, cell differentiation and proliferation ^9^. NF-κB activation involves the degradation of IκB that sequesters NF-κB factors in the cytoplasmic compartment, leading to NF-κB nuclear translocation and binding of NF-κB dimers (e.g., RELA:p50 for the most abundant) to their target promoters ^10,11^. TAX induces IKK phosphorylation and IκB degradation, leading to persistent nuclear translocation of NF-κB ^12,13^. In addition, TAX interacts with nuclear NF-κB factors and enhances their effects on transcription ^14,15^.

Interestingly, genome-wide analyses of NF-κB distribution have unveiled that the vast majority of RELA peaks is outside promoter regions and can be localized in introns and exons ^16–19^. Some of those promoter-distant RELA binding sites correspond to *cis*-regulatory transcriptional elements ^20,21^ but globally, there is a weak correlation between the binding of RELA to genes and regulation of their steady-state expression ^17,18^. These data suggest that NF-κB could have other functions than its initially described transcription factor function.

Here, we show for the first time that NF-κB activation accounts for alternative splicing modifications generated upon TAX expression. These effects rely on a tight physical and functional interplay between TAX, RELA and the DDX17 splicing factor. Our results reveal that DNA binding of RELA at the vicinity of genomic exons regulates alternative splicing through the recruitment of DDX17, which modulates exon inclusion thanks to its RNA helicase activity.

## Results

### TAX induces alternative splicing modifications irrespectively of its effects on transcription

RNA-seq analyses were performed on 293T-LTR-GFP cells transiently transfected with a TAX expression vector. TAX-induced changes in gene expression level and in alternative splicing were identified and annotated as previously described ^22,23^(Table S1). As shown in Figure 1A, the ectopic expression of TAX affected the splicing and gene expression levels of 939 and 523 genes, respectively. A total of 1108 alternative splicing events were predicted including 710 exon skipping events (Figure 1B). A minority of genes (3.5%, 33/939) was altered at both the expression and splicing levels, indicating that TAX largely affects alternative splicing independently of its transcriptional activity. A subset of splicing events was validated by RT-PCR (Figure 1C). We took advantage of RNA-seq datasets (EGAS00001001296 ^24^) for assessing whether TAX-related alternative splicing could pertain to asymptomatic carriers (AC) and ATLL patients. Overall, 542 (48%) TAX-induced splicing modifications were detected at least once across 55 clinical samples (Table S1). Hierarchical clustering of these exons based on their inclusion rate (PSI) identified TAX-regulated exons that discriminate AC and ATLL samples from uninfected CD4+ T-cells (Figure 1D). We furthermore confirmed that TAX promotes splicing events previously detected in HTLV-1 infected individuals, including *AASS*, *CASK*, *RFX2* and *CD44* ^4,25^. We firmly established that the expression of the splicing variant *CD44v10* previously identified in HAM/TSP patients ^25^ fully relies on TAX expression (Figure 1C and Figures S1A-C). Altogether, these results uncovered a large number of splicing modifications upon TAX expression that for a part coincide with alternative splicing events observed in HTLV-1 patients.

**Figure 1:**
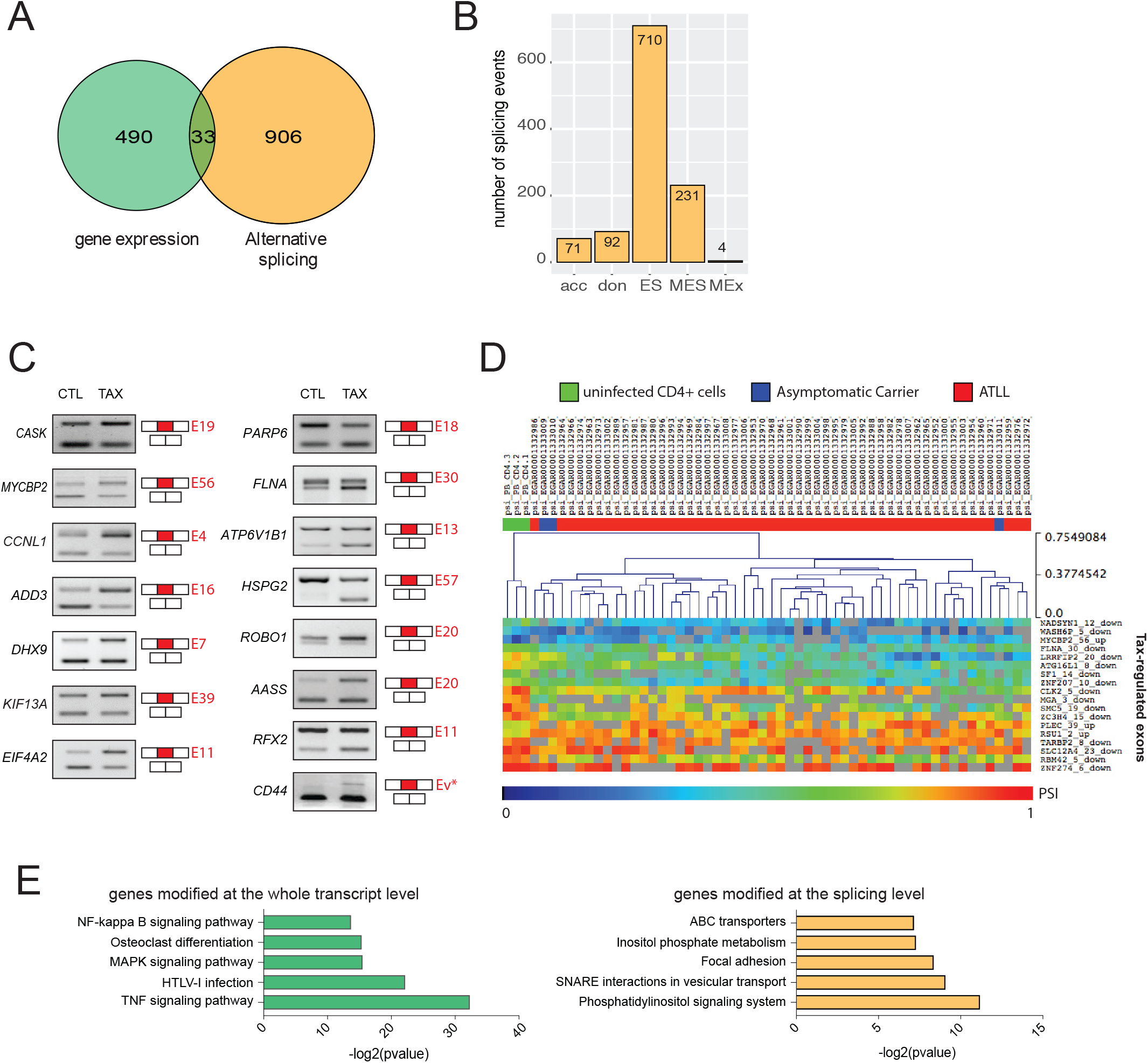
TAX induces alternative splicing modifications independently of its transcriptional effects. (A) Genes regulated at the steady-state expression level and at the splicing level upon TAX expression. The significance thresholds were typically set to 10% for ∆PSI (differential percentage of spliced-in sequence) and 0.6 for log2-gene expression changes (p-val<0.05, Fisher’s exact test), respectively. (B) Different alternative splicing events induced by TAX: alternative acceptor (acc), alternative donor (don), exon skipping (ES), multi-exon skipping (MES), multi-exclusive exon skipping (MEx). (C) Validation of alternative splicing predictions by RT-PCR (35 cycles of PCR). The exon number is indicated in red. CD44 full variants (Ev*) were assessed using primers C13 and C12A (Figure S4).(D) Exon-based hierarchical clustering. Kruskal-Wallis ANOVA (p-val<0.05) was carried out with Mev4.0 software (http://www.tm4.org/) using the PSI values of exons that share similar regulations upon TAX and in clinical samples (EGAS00001001296). Only the most significant exon regulations are presented. (E) Gene ontology analysis (DAVID) of TAX splicing and transcriptional targets.

Gene ontology analysis of quantitatively altered genes revealed several signaling pathways that are well described in TAX expressing cells, including NF-κB, TNF, and MAPK signaling (Figure 1E) ^26,27^. In contrast, genes modified at the splicing level belong to membrane-related regulatory processes including focal adhesion and ABC transporters (Figure 1E). In this setting, we observed that TAX-expressing cells displayed switched cell adhesion properties from hyaluronate-to type IV collagen-coated surfaces, which is in accordance with the substrate affinity of the CD44v10 isoform ^28^ (Figure S1D).

### The splicing factor DDX17 interacts with RELA and TAX in a NF-κB dependent manner

Since Tax is a well-known trans-acting transcription regulator, we first analysed whether TAX could affect gene expression levels of splicing factors. However, no significant change was measured for 227 genes encoding splicing regulators (Table S1, Figure 2A), thereby suggesting a direct role of TAX in alternative splicing regulatory mechanisms. To tackle this question, we focused on the auxiliary component of the spliceosome DDX17, which has been previously identified, but not validated, in a recent mass spectrometry screen for putative protein partners of TAX ^29^.

**Figure 2:**
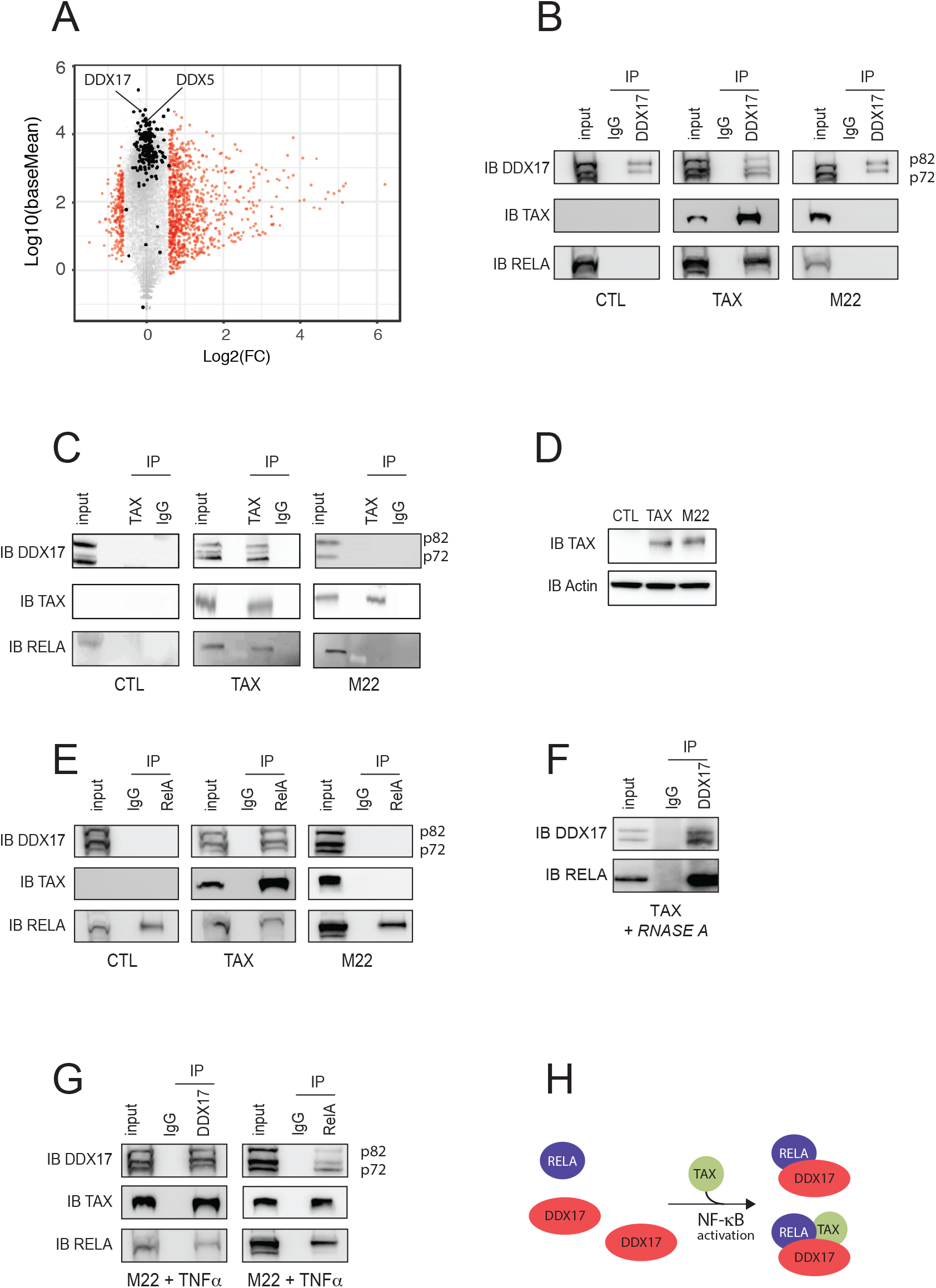
Physical interactions between TAX, RELA and DDX17 in an NF-κB dependent manner. Mean average plot (n=3, p<0.05) of cellular gene expressions upon TAX. Each gene is plotted according to its expression level (Log10(BaseMean) from DESeq2 analysis) and to fold change (Log2-FC) upon TAX. Red dots show significant gene expression changes (Log2-FC>0.6, p-val<0.05, Fisher’s exact test). Black dots highlight genes encoding splicing factors. DDX5 and DDX17 are indicated. (B) Immunoprecipitation assays (IP) were carried out using isotype IgG or anti-DDX17 (B and G), anti-RELA (E and G) and anti-TAX (C and G) antibodies, followed by immunoblotting (IB) with indicated antibodies. (D) Western blot analysis of TAX and M22 expression 48h post-transfection. (F) RNA-free IP assays. (G) TNFα exposure of M22 expressing cells promotes RELA-DDX17 interactions. (H) Model of NF-κB-dependent interplay between TAX, RELA and DDX17.

We therefore aimed to validate the interaction between TAX and DDX17. As shown in Figure 2B, TAX co-immunoprecipitated with the two endogenous isoforms of DDX17, namely p72 and p82. Reciprocal IP confirmed this interaction (Figure 2C). Due to the involvement of NF-κB signaling in TAX positive cells (Figure 1D, ^27^), we examined whether DDX17 interacts with a TAX mutated form, namely M22 (G137A, L138S), which is defective for IKK and NF-κB activation ^30–33^. Despite similar expression levels and immunoprecipitation efficiencies of TAX and M22 (Figure 2D), we failed to detect any interaction between M22 and DDX17 (Figures 2B and 2C), suggesting that NF-κB is required for recruiting DDX17. In this setting, RELA co-immunoprecipitated with DDX17 and TAX, but not with M22 (Figures 2B and 2C). Moreover, DDX17 was co-immunoprecipitated with RELA in a TAX-dependent manner (Figure 2E). This interaction did not require RNA since the DDX17:RELA complex remained detected when cell extracts were pre-treated with RNAse A (Figure 2F).

As DDX17:RELA complexes were observed neither in control cells (that do not expressed TAX) nor in M22 expressing cells, this suggested that NF-κB activation is necessary for the binding of DDX17 to RELA. This hypothesis was confirmed by exposing TAXM22-expressing cells to TNFα, a potent NF-κB activator that allowed to retrieve DDX17:RELA complexes (Figure 2G). Altogether, these results revealed that TAX-induced NF-κB activation dynamically orchestrates the interations between TAX, the transcription factor RELA and the splicing regulator DDX17 (Figure 2H).

### TAX-mediated effects on splicing depend on DDX5/17

To estimate the role of DDX17 in TAX-regulated splicing events, RNA-sequencing was performed using 293T-LTR-GFP cells expressing or not TAX and depleted or not for DDX17 and its paralog DDX5, which cross-regulate and complement each other ^22, 34, 35^. TAX had no effect on the expression of DDX5 and DDX17 (Figures 2A and 3A) and RELA protein level was not significantly changed upon both TAX expression and *DDX5/17* silencing (Figure 3B).

**Figure 3:**
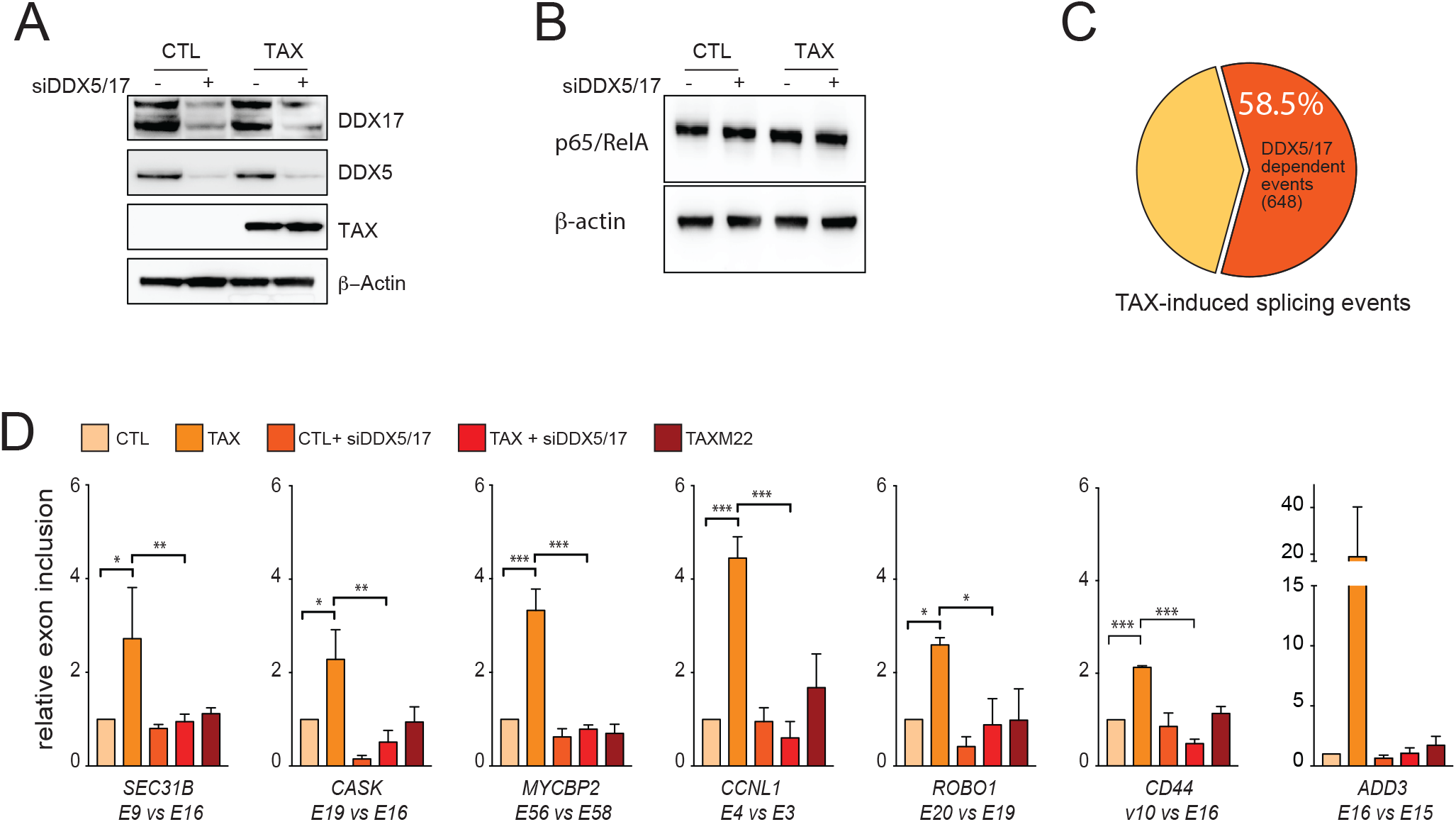
DDX5/17 regulates TAX splicing targets in an NF-κB-dependent manner. (A) Western blot analysis of DXX5 and DDX17 expression in cells expressing or not TAX and depleted or not of DDX5 and DDX17 by siRNA. (B) Western blot analysis of RELA and β-actin upon TAX expression and siRNA-DDX5/17 delivery. (C) Splicing events modified upon the depletion of DDX5/17 in TAX positive cells. The significant threshold was set to ≥2 in comparisons between TAXvsCTL and TAXsiDDX5/17vsCTL. (D) Validation of alternative splicing predictions of a set of TAX- and DDX5/17-regulated exons. Histograms represent the results of exon specific quantitative RT-PCR measurements computed as a relative exon inclusion (alternatively spliced exon vs constitutive exon reflecting the total gene expression level) from three biological replicates ± s.d.. All of these genes but MYCBP2 were unmodified at the whole transcript level upon TAX expression (Figure S2C).

Overall, 58.5% (648/1108) of TAX-regulated exons were affected by *DDX5/17* knockdown, a significantly higher proportion than expected by chance (Figure 3C, Figure S2A). Of particular significance, 423 TAX-induced splicing events were completely dependent on the presence of DDX5/17 (Table S3). For example, *DDX5/17* silencing completely abolished the TAX-mediated effect on splicing of *SEC31B*, *CASK*, *MYCBP2*, *CCNL1*, *ROBO1, ADD3* and *CD44* transcripts (Figure 3D). Of note, splicing specific RT-PCR assays permitted to validate the effect of DDX5/17 on TAX-dependent splicing changes for *CD44*, *ADD3* and *EIF4A2* transcripts, even though their predicted differential inclusion fell below the arbitrary computational threshold (Table S3, Figure 3D and Figure S2D). This suggested that the contribution of DDX5/17 to TAX-mediated alternative splicing regulation might be under-estimated.

Finally, since NF-κB activation modified the interactions between DDX17, RELA, and TAX (Figure 2), we examined the interplay between NF-κB activation and DDX17-mediated splicing regulation. As shown in Figure 3D, M22 did not have any effect on DDX5/17-sensitive splicing events, arguing that TAX splicing targets are regulated by RNA helicases DDX5/17 in an NF-κB dependent manner.

### RELA binds to genomic exons and recruits DDX17 to regulate splicing in an RNA helicase-dependent manner

The results described above prompted us to hypothesize that the nuclear translocation of RELA upon TAX expression might promote the chromatin recruitment of DDX17 to RELA target genes. To test this hypothesis, the *CD44* gene was used as a gene model. *CD44* is composed of 10 constitutive exons and 10 variable exons. The constitutive exons 1–5 and 15–20 encode the standard *CD44* transcripts, while *CD44* variants (*CD44v*) are produced by extensive splicing leading to alternative inclusion of variable exons 5a-14 also named v1-v10 (Figure 4A) ^36^. As shown above (Figure 3D), the exon v10 inclusion rate is markedly influenced by TAX in a DDX5/17- and NF-κB activation-dependent manner. The importance of NF-κB in this process was further confirmed as the inactivation of NF-κB *via* the ectopic expression of the IkBα super repressor (IkBSR) abolished the effects of TAX on *CD44* v10 inclusion (Figure S3A).

**Figure 4:**
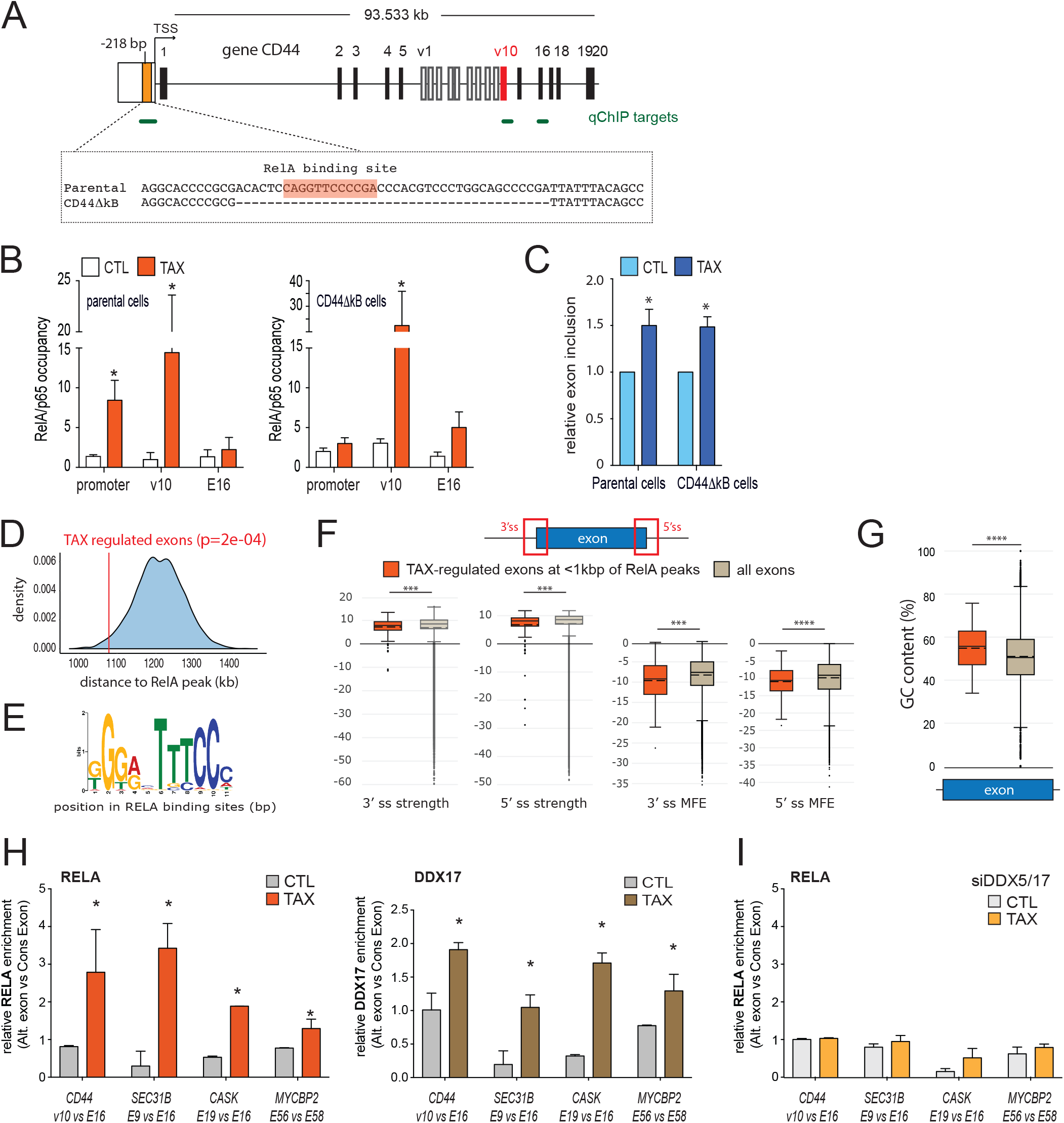
RELA locally recruits DDX17 at the genomic target exons for regulating splicing. (A) Schematic representation of the human CD44 gene. Black and white boxes represent constitutive and alternative exons as previously annotated (50), respectively. The orange box represents the κB site localized at −218 bp from the TSS and the 40 bp fragment deleted by CRISPR/Cas9 in CD44ΔkB cells. (B) qChIP analysis of RELA occupancy across the promoter, the exon v10, and the constitutive exon E16 of CD44. The RELA enrichment is expressed as the fold increase in signal relative to the background signal obtained using a control IgG. (C) Relative exon inclusion of CD44 exon v10 were quantified by qRT-PCR in parental cells and its CD44∆kB counterparts. The histogram shows mean ± sd of three independent experiments. (D) Bootstrapped distribution of median distance between intragenic RELA peaks and either TAX-regulated exons (red line, 1079 bp) or randomly chosen exons (10000 repetitions) (blue). pval: sample t-test (E) Consensus de novo motif for RELA binding sites <1kbp of TAX regulated exons. (F) Strength and minimum free energy (MFE) of 3’ and 5’ splice sites, and GC content (G) of TAX-regulated exons localized at <1kbp of RELA binding site. The exons of FasterDB database were used as control. (H) Relative RELA and DDX17 occupancies at TAX-regulated exons in control and TAX expressing cells. For each gene, the binding of RELA or DDX17 to the regulated exon is represented as a comparison to the binding to a neighbouring constitutive exon (*p-val≤0.05). (I) Relative RELA occupancy at TAX-regulated exons in cells treated with DDX5/17-specific siRNA and expressing or not TAX. Details are as in D. (*p≤0.05,***p≤0.001, ****p≤0.0001, Mann-Whitney test)

Quantitative ChIP (qChIP) analyses revealed that RELA was recruited upon TAX expression not only to the *CD44* promoter, but also to a genomic region spanning the alternative exon v10, but not a downstream constitutive exon (E16) (Figures 4A and 4B, left panel). To assess whether RELA occupancies at the v10 exon and CD44 promoter are interrelated, a stable cell line was generated in which the κB site localized at −218 bp from the transcription start site (TSS) was deleted using a CRISPR-Cas9 approach. Positive clones (CD44ΔkB) were screened and sequenced to confirm the 40 bp deletion in the promoter region (Figure 4A). As expected, TAX expression failed to promote RELA binding at the promoter in CD44ΔkB cells (Figure 4B, right panel). Nevertheless, TAX still promoted RELA binding at the v10 region. Importantly, TAX expression induced v10 inclusion at a similar level in both CD44ΔkB and parental cells (Figure 4C). These results suggested that TAX-mediated effect on exon v10 splicing could depend on RELA binding in the vicinity of the alternative v10 exon. Supporting this hypothesis, the analysis of publicly available RELA ChIP-seq datasets revealed that intragenic RELA peaks are significantly closer to alternative exons than to constitutive exons (Figure S3B). In this setting, we observed that RELA binding sites are often found in the vicinity of TAX regulated exons (Figure 4D). Using the MEME-ChiP suite as motif discovery algorithm^37^, we uncovered that RELA-binding sites located within the closest range (<1kbp) of TAX-regulated exons coincided with the typical NF-κB consensus motif (Figure 4E). Furthermore, this subset of TAX-regulated exons displayed weak 3’ and 5’ splice sites together with significant low MFE value (Figure 4F) and high GC-content (Figure 4G) when compared to all human exons. This emphasizes the high potential of these splice sites to form stable secondary RNA structures, a typical feature of exons regulated by RNA helicases DDX5/17 ^34^. Taken together, these data define a signature of splicing target specificity for RELA, and they suggest that RELA and DDX17 might control together the inclusion of a subset TAX-regulated exons. We therefore investigated the genomic occupancy of some target exons by RELA and DDX17 by qChIP analysis of cells expressing or not TAX. For all tested genes (*CD44*, *SEC31B*, *CASK*, and *MYCBP2*), both RELA and DDX17 bound specifically the regulated alternative exon in a TAX-dependent manner, compared to a downstream constitutive exon (Figure 4H). Furthermore, RELA binding was lost in cells depleted for DDX5/17, indicating that RNA helicases contribute to stabilize DNA-bound RELA (Figure 4I).

### A causal relationship between exonic DNA-binding of RELA, chromatin recruitment of DDX17, and splicing regulation

To assess the causative relationship that links RELA and DDX17 to alternative splicing, we intended to experimentally tether DDX17 or RELA at the *CD44* v10 exon locus using modified TALE (Transcription-Activator-Like-Effector) ^38^. We designed a TALE domain that recognizes specifically an exonic 20 bp DNA sequence located 12 bp upstream from the 5′ splice site (SS) of exon v10. This TALE domain was either fused to RELA or DDX17 proteins (Figure 5). We also used an additional construct consisting in the same TALE fused to GFP to rule out non-specific effects resulting from the DNA binding of the TALE. Each TALE construct was transiently transfected into 293T-LTR-GFP cells, and we monitored their relative effects both on the recruitment of endogenous RELA and DDX17, and on the splicing of exon v10. All results shown in Figure 5 were normalized and expressed as relative effects compared to the TALE-GFP. As expected, and validating our approach, TALE-RELA tethering to the exon v10 led to a significant chromatin recruitment of RELA to its target site, and not to the downstream exon E16 used as control (Figure 5A, left panel). A significant and specific enrichment of DDX17 was also observed on exon v10 upon expression of the TALE-RELA compared to TALE-GFP (Figure 5A, left panel), indicating that tethering RELA to exon v10 induced a local recruitment of endogenous DDX17 proteins. At the RNA level, this TALE-RELA-mediated recruitment of DDX17 coincided with a significant increase in exon v10 inclusion rate (Figure 5A).

**Figure 5:**
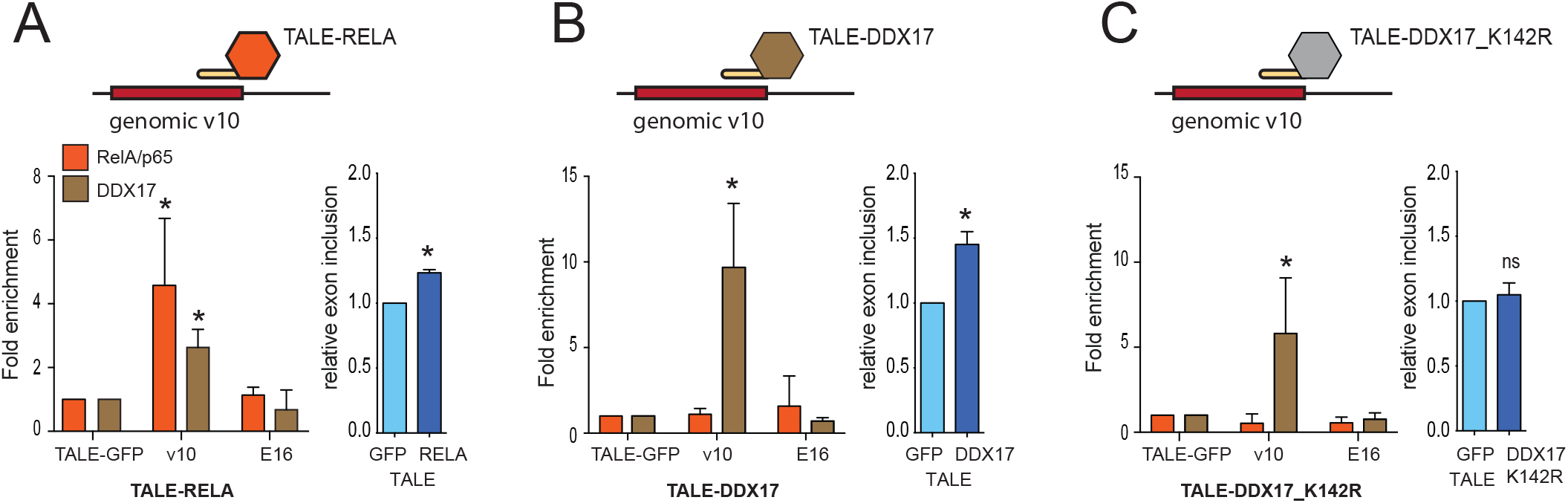
Chromatin and splicing regulation upon TALE-mediated tethering of RELA and DDX17. (A) The TALE domain was designed to bind the v10 exon of CD44 and fused to either GFP (A-C), RELA (A), DDX17 (B) or its helicase-deficient mutant DDX17_K142R (C). The effect of TALEs on RELA and DDX17 chromatin enrichment (left panels) and on the relative v10 exon inclusion (right panels) was monitored by qChIP and qRT-PCR, respectively. Results were normalized to measures obtained in TALE-GFP assays. Mean ± sd of three independent experiments are shown (*p-val<0.05).

We next investigated whether DDX17 tethering could result in similar effects. Quantitative ChIP analysis demonstrated that DDX17 was properly tethered to exon v10 when fused to the designed TALE (Figure 5B) but TALE-DDX17 had no effect on RELA recruitment (Figure 5B). This was expected since the formation of RELA:DDX17 complexes only occurs upon NF-κB activation (Figure 2). Nevertheless, TALE-DDX17-expressing cells exhibited a reproducible and significant increase in v10 inclusion (Figure 5B), indicating that chromatin-bound DDX17 alone can modulate splicing efficiency. It is worth to underline that the level of v10 exon inclusion induced by the TALE-RELA and - DDX17 was comparable to that measured in cells transiently transfected with a TAX expression vector (Figure 4C and Figure S1C). Although it is less quantitative approach, a nested RT-PCR assay clearly confirmed these results (Figure S4). Strikingly however, the TALE-DDX17_K142R (a DDX17 helicase mutant ^34,39-41^) failed to influence exon v10 inclusion despite a clear chromatin enrichment of DDX17 (Figure 5C, Figure S4). Collectively, these results demonstrate that the binding of RELA at the vicinity of genomic exons recruits the RNA helicase DDX17 that positively regulates the inclusion rate of the target exon thanks to its RNA helicase activity.

## DISCUSSION

Since the finding of splicing dysregulations in HTLV-1 infected individuals ^4,24,25,42^, deciphering how HTLV-1 interferes with the splicing regulatory network has become a new challenging issue for improving our knowledge of HTLV-1 infection and its associated diseases. Here, we provide the first molecular evidence that upon TAX-induced NF-κB activation, RELA directly regulates splicing by binding to gene bodies at the vicinity of GC-rich exons and by locally recruiting the splicing factor DDX17, which regulates splicing via its RNA helicase activity.

Our results demonstrate for the first time that TAX deeply impacts alternative splicing independently from its effects on transcription. In addition, TAX-regulated exons were found in transcripts enriched in functional pathways that are distinct from those enriched by TAX transcriptional targets, suggesting that splicing reprogramming may constitute an additional layer of regulations by which HTLV-1 modifies the host cell phenotype. Arguing for this, we showed that the TAX-induced splicing variant *CD44v10*, which was previously identified in circulating blood of HAM/TSP patients ^25^ and confirmed here *ex vivo* in infected CD4+ T-cell clones, contributes to modulate cell adhesion affinity *in vitro*. GO analyses of TAX splicing targets also pointed to the phosphatidylinositol signaling system and to the inositol phosphate metabolism, two processes that are particularly connected to NF-κB signaling and that play critical roles in oncogenesis and disease progression of malignant diseases, including ATLL ^43,44^. This suggests that, besides its transcriptional effects, splicing regulatory functions of TAX might account for its oncogenic properties. Accordingly, a large number of TAX-regulated exons could be observed in ATLL samples, which rarely express TAX but typically exhibit NF-κB addiction for survival and proliferation ^24, 45, 46^.

At the molecular level, we showed that the increased chromatin occupancy of RELA upon TAX expression is not restricted to promoter regions but also occurs in the vicinity of exons that are regulated at the splicing level (Figure 4). Exons regulated by TAX, especially those localized within 1 kb of intragenic RELA binding sites, are characterized by a high GC-content, a typical feature of exons regulated by the DDX5 and DDX17 RNA helicases ^34^ (Figure S3D). Accordingly, we found that a majority of TAX-regulated exons depend on the expression of these proteins (Figure 3). A local chromatin recruitment of DDX17 and RELA was validated on several TAX-regulated exons (Figure 4). More importantly we identified a confident causal relationship between the exonic tethering of RELA, the local chromatin recruitment of DDX17, and the subsequent splicing regulation via DDX17 RNA helicase activity. This catalytic activity of DDX17 is strictly required for its splicing regulatory functions (Figure 5), as previously reported ^34^. Indeed, the RNA helicase activity of DDX5 and DDX17 has been involved in resolving RNA structures, facilitating the recognition of the 5’ splice site, that can be embedded in secondary structures, and exposing RNA binding motifs to additional splicing regulators ^34,40,47–49^. However, even though some RNA binding specificity has been reported for DDX17 ^50,51^, these RNA helicases are devoid of a proper RNA binding domain and their activity in splicing may depend on additional factors that are able to provide target specificity. Here, we suggest that RELA may be also regarded as a DDX17 recruiter by acting as a chromatin anchor for DDX17 in the vicinity of exons dynamically selected upon NF-κB activation.

The target specificity of NF-κB factors remains a complex question. It has been estimated that approximately 30 to 50% of genomic RELA binding sites do not harbor a typical NF-κB site, and that only a minority of RELA-binding events associate with transcriptional change^16–19^, thereby indicating that neither a consensus site nor significant NF-κB occupancy are sufficient criteria for defining RELA’s target specificity. Here, we identified a typical κB consensus motif at RELA-binding loci that are close to alternatively spliced exons but we also uncovered that weak splice sites, low MFE, and significant GC-content bias of exons likely contribute to RELA’s target specificity. Because low MFE and high GC-content confer a high propensity to form stable RNA secondary structures, the recognition and the selection of such GC-rich exons with weak splice sites by the splicing machinery typically depend on RNA helicases DDX5/17 ^34^. Based on these observations we propose that, upon TAX-induced NF-κB activation, RELA binds to intragenic binding consensus motifs and locally recruits DDX17. When the RELA:DDX17 complex is located at a close proximity of GC-rich exons flanked by weak splice sites, DDX17 can impact on their inclusion rate by unwinding GC-rich secondary structures of the nascent RNA transcript, and by potentially unmasking binding motifs for additional splicing regulators.

In conclusion, our results provide conceptual advance for understanding how cell signaling pathways may drive target specificity in splicing by dynamically recruiting cognate transcription factors at the vicinity of target exons that act as chromatin anchor for splicing regulators. In the context of NF-κB signaling, such mechanism likely has a significant impact on cell fate determination and disease development associated with HTLV-1 infection, but also on other situations linked to chronic NF-κB activation, as numerous human inflammatory diseases and cancer.

## Acknowledgments

This study makes use of data (EGAS00001001296) generated by Department of Pathology and Tumor Biology, Kyoto University ^24^. We gratefully acknowledge support from the PSMN (Pôle Scientifique de Modélisation Numérique) of the ENS de Lyon for the computing resources. This work was supported by the Ligue Contre le Cancer (Comité de la Savoie, de la Drome et du Rhône), the Fondation ARC (ARC PJA20151203399), and the Agence Nationale pour la Recherche (program EPIVIR and CHROTOPAS). L.B.A was supported by Ligue Contre le Cancer. G.G was support by ANR CHROTOPAS. M.T. was supported by a bursary from the French Ministry of Higher Education and Science. F.M., C.F.B, and D.A. were supported by INSERM. E.W. was supported by Hospices Civils de Lyon and Lyon I University (France).

## Author Contributions

Conceptualization, L.B.A., D.A. and F.M.; Resources, A.G. and E.W.; Experiments, L.B.A., M.T., and G.G.; Technical Support, E.C. and M.B.; Bioinformatic, J.B.C., H.P. and N.F., Formal Analysis, L.B.A., M.T., G.G. and F.M.; Supervision, D.A. and F.M.; Funding Acquisition, C.F.B., D.A. and F.M.; Writing – Original Draft, F.M.; Writing – Review and Editing, L.B.A, M.T., G.G., C.F.B., D.A. and F.M.

## Supplementary data

**Table S1: Whole gene expression and alternative splicing changes upon TAX expression.** Whole gene expression threshold was set to log2FC=0.6, p<0.05. For differential splicing, deltaPsi threshold was set to 1.1 p<0.05 (Fisher test). TAX-induced splicing regulations identified in RNA-seq datasets derived from carriers and ATLL samples (EGAS00001001296). Alternative splicing profiles of each clinical sample was assessed using Farline analysis with peripheral blood CD4+ T-cells used as control. The table lists 452 TAX-induced splicing regulations identified at least once across 56 clinical samples.

**Table S2: sequence features of exons regulated by TAX**

**Table S3: genes modified in splicing by TAX in a DDX5/17-dependent manner.** TAX splicing targets responsive to DDX5/17 were attributed to splicing events lost upon DDX5/17 depletion.

**Table S4: list of oligonucleotides, siRNA, sgRNA and TALE sequences**

## Methods

### Cell Culture and Transfections

Peripheral blood mononuclear cells (PBMCs) were obtained by Ficoll separation of whole blood of HTLV-1 infected individuals. T-cell limiting dilution cloning was performed as previously described ^4^. The human embryonic kidney 293T-LTR-GFP cells ^52^, which contain an integrated GFP reporter gene under the control of the TAX-responsive HTLV-1 LTR, were cultured in DMEM+Glutamax medium supplemented with decomplemented 10% FBS and 1% penicilline/streptomycine. This cell line was used to measure transfection efficiency in TAX and TAX M22 conditions. In standard transfection experiments, siRNAs (Table S4) and/or expression vectors (pSG5M empty, pSG5M-TAX-WT, pSG5M-M22) were mixed with JetPrime (Polyplus-Transfection) following the manufacturer’s instructions and cells were harvested 48h after transfection. TNFα exposure consisted in treating cells with 10ng/ml of TNFα for 24 hours.

### Cell-adhesion assays

Culture plates were prepared by coating with 40 µg/ml hyaluronic acid from human umbilical cord (Sigma) and 25 µg/ml type IV collagen from human placenta (Sigma) overnight at 4°C. Non-specific binding sites were blocked for 1 h with PBS containing 1 mg/ml heat-denatured BSA. After three washes with PBS, 5 × 10^4^ cells transiently transfected with pSG5M-TAX vector or its empty control were added at 48h post-transfection. Cell adhesion was allowed to proceed for 20 min at room temperature. Non-adherent cells were removed with 3 PBS washes, and adherent cells were quantified. All experiments were done in triplicate.

### Western Blot

Cells were washed twice with 1X PBS and total proteins were directly extracted in RIPA buffer (50mM Tris HCL pH 7.4, 50 mM NaCl, 2mM EDTA, 0.1% SDS). A total of 20 μg of whole cell proteins were separated on a NuPAGE™ 4-12% Bis-Tris Protein Gels and transferred on a nitrocellulose membrane using Trans-Blot® Turbo^TM^ Blotting System. Membranes were saturated with 5% milk and incubated overnight at 4°C with the primary antibodies against RELA (sc-109, Santa Cruz), Tax (1A3, Covalab), DDX17 (ProteinTech), DDX5 (ab10261 Abcam), Actin (sc-1616, SantaCruz). After three washes with 1x TBS-Tween, membranes were incubated 1h at room temperature with the secondary antibodies conjugated with the HRP enzyme and washed 3 times as above. Finally, the HRP substrate (GE Helathcare) was applied to the membrane for 5 minutes and the chemiluminescence was read on Chemidoc (Biorad).

### Co-immunoprecipitation

Cells were harvested in IP lysis buffer (20 mM Tris-HCl pH 7.5, 150 mM NaCl, 2 mM EDTA, 1% NP40, 10% Glycerol). Extracts were incubated overnight with 5 µg of antibodies recognizing either RELA (C20 sc-372, Santa Cruz), Tax (1A3, Covalab), DDX17 (ProteinTech) in the presence of 30 μL Dynabeads® Protein A/G (Thermo Fisher). Isotype IgG rabbit (Invitrogen) or mouse (Santa Cruz) was also used as negative control. The immunoprecipitated complexes were washed three times with IP lysis buffer.

### Chromatin Immunoprecipitation

A total of 10^7^ cells were crosslinked with 1% formaldehyde for 10 minutes at room temperature. Crosslinking was quenched by addition of 0.125 M glycin. Nuclei were isolated by sonication using a Covaris S220 (2 min, Peak Power: 75; Duty Factor: 2; Cycles/burst: 200), pelleted by centrifugation at 3,000 rpm for 5 min at 4°C, washed once with FL buffer (5 mM HEPES pH 8.0, 85 mM KCl, 0,5% NP40) and resuspended in 1 mL shearing buffer (10 mM Tris-HCl pH 8.0, 1 mM EDTA, 2 mM EDTA, 0,1% SDS). Chromatin was sheared in order to obtain fragments ranging from 200 to 800 bp using Covaris S220 (20 min, Peak Power: 140; Duty Factor: 5; Cycles/burst: 200). Chromatin was next immunoprecipitated overnight at 4°C with 5 µg of antibodies: RELA (C20 sc-372, Santa Cruz), DDX17 (19910-1-AP, ProteinTech) and V5 (AB3792, Millipore). Then, 30 μL Dynabeads® Protein A/G (Thermo Fisher) were added. Complexes were washed using 5 different buffers: Wash 1 (1 % Trition, 0.1 % NaDOC, 150mM NaCl, 10mM Tris HCL pH8), Wash 2 (1 % NP-40, 1 % NaDOC, 150mM KCl, 10mM Tris HCL pH8), Wash 3 (0.5 % Trition, 0.1 % NaDOC, 500mM NaCl, 10mM Tris HCL pH8), Wash 4 (0,5% NP-40, 0.5 % NaDOC, 250mM LiCl, 20mM TRIS.Cl pH8, 1mM EDTA), Wash 5 (0.1 % NP-40, 150mM NaCl, 20mM Tris HCL pH8,1mM EDTA). The immunoprecipitated chromatin was purified by phenol-chloroform extraction and quantitative PCR was performed using Rotor-Gene 3000 cycler (Corbett) or LightCycler 480 II (Roche, Mannheim, Germany). Values were expressed relative to the signal obtained for the immunoprecipitation with control IgG. Primers used for ChIP experiments were designed in exon/intron junction (Table S4). For TALE ChIP experiment, DDX17 and RelA enrichment were normalized to the signal observed with V5 antibody corresponding to TALE recruitment. Then, the TALE GFP condition was used as control and set to 1.

### RNA extraction, classical PCR and Real-time quantitative PCR

Total RNAs were extracted using TRIzol (Invitrogen). RNAs (2.5 μg) were retro-transcribed with Maxima First Strand cDNA Synthesis Kit after treatment with dsDNase (Thermo Scientific) following the manufacturer’s instructions. PCRs were performed using 7.5 ng of cDNAs with GoTaq polymerase (Promega, Madison, WI, USA). PCR products were separated by ethidium bromide-labeled agarose gel electrophoresis. Band intensity was quantified using the ImageLab software (Bio-Rad). Quantitative PCR was then performed using 5 ng of cDNAs with SYBR® Premix Ex Taq TM II (Tli RNaseH Plus) on LightCycler 480 II. Relative level of the target sequence was normalized using the 18S or GAPDH gene expression (∆Ct) and controls were set to 1(∆∆Ct). We calculated the inclusion rate of alternative exons using the following method: 2^−∆∆Ct^ (included exon)/2^−∆∆Ct^ (constitutive exon). The oligonucleotide sequences used are listed in Table S4.

### RNA-seq and bio-informatic analysis

RNA-seq analyses were performed as previously described ^22^. Briefly, poly-A transcripts were extracted from 293T-LTR-GFP cells transfected with pSG5M-Tax or pSG5M empty vectors and knockdown or not for DDX5-17. RNA-seq libraries were generated at the Aros Applied Biotechnology (Aarhus, Denmark) using Stranded mRNA Sample Prep kit (Illumina) and sequenced using the illumina HiSeq 2500 technology. Each sample have in average 6.10^7^ of paired-end pairs of reads. These RNA-seq data were analyzed using FaRLine, a computational program dedicated to analyzing alternative splicing with FasterDB database ^23,53^. The gene expression level in each sample was calculated with HTSeq-count (v0.7.2) ^54^ and differential expression between conditions was computed with DESeq2 (v1.10.1) (abs(log2FoldChange) ≥ 0.4, *p*values ≤ 0.05) ^55^. In silico screening of NF-κB responsive elements in the CD44 promoter sequence was carried out via PROMO database (based on TRANSFAC v8.3) ^56^. The MEME-ChIP suite was used to discover the regulatory motifs in the NF-kB ChIP-seq data ^37^.

For the prediction of splice site strength, scores were computed using MaxEntScan ^57^ for sequence (3 bases in the exon and 6 bases in the intron for 5’ splice sites; 20 bases in the intron and 3 bases in the exon for 3’splice sites) covering both sides of the splicing site. MaxEntScan uses Maximum Entropy Models (MEMs) to compute log odds ratios. The minimum free energy was computed from exon-intron junction sequences using RNAFold from the ViennaRNA package (v 2.4.1; http://rna.tbi.univie.ac.at/cgi-bin/RNAWebSuite/RNAfold.cgi). Analyzed sequences include 25 nucleotides within the intron and 25 nucleotides within the exon. The GC content was calculated for exons defined in FasterDB ^53^.

The distribution of RELA peaks across alternative and constitutive exons, and the average distance between RELA peaks and TAX exon targets was measured using ChiP-seq datasets from GEO ^58^, ENCODE ^59^ and CISTROME ^60^ databases: from GEO GSE63736, GSM1239484, GSM486271, GSM486293, GSM486298, GSM486318, GSM847876, GSM847877, GSM2394419, GSM2394421, GSM2394423, from ENCODE ENCFF002CPA, ENCFF002CQB, ENCFF002CQJ, ENCFF002CQN, ENCFF580QGA and from CISTROME 53597, 5388, 5389, 4940, 36310, 36316, 4971. For another GEO dataset, GSM2628088, reads were mapped to the hg19 build of the human genome with Bowtie2 ^61^ and RelA peaks were identified with Macs2 ^62^. Alternative and Constitutive spliced exons were obtained from FasterDB ^53^. In order to focus on intragenic RELA peaks, we used the bedtools ^63^ intersect command to remove all intergenic RELA peaks and all RELA peaks localized on first exon (or at least at less than 500nt) for each gene. A Perl script was specifically created to measure the distance between RELA peaks and TAX-regulated exons. Briefly, RELA peaks and exons are provided as BED files and the script reports for each exon the distance in nucleotides of the nearest RELA peak. Closest peak distances from the 710 TAX-regulated exon-cassettes were compared to closest peak distances from 710 exons chosen by chance (10^5^runs).

### TALE design and construct

The TALE constructs were obtained from ThermoFisher Scientific. TALEs were constructed using the Golden Gate Assembly method as previously described ^38^. The RVDs HD, NI, NG and NN were chosen to specifically recognize the nucleotides C, A, T and G, respectively. The TALE targeting CD44 v10 sequence was 5’ TCCAACTCTAATGTCAATC 3’. This TALE construct was fused to a V5 sequence and a SV40 NLS at its 5’ end and cloned in the NotI-HindIII fragment of the pXJ41 backbone plasmid. DDX17-WT and DDX17-K142R cDNA were obtained by PCR from pcDNA3-HA-DDX17 and pcDNA3-HA-DDX17-K142R and were cloned in the HindIII-BglII fragment in the MCS downstream to the TALE sequence. The *RELA* cDNA was amplified from a library of cDNA of 293T-LTR-GFP cells and was cloned in the HindIII-BamHI fragment.

### CRISPR design and construct

The sequence-specific sgRNA for site-specific interference of genomic targets were designed using CRISPRseek R package^1^, and sequences were selected to minimize off-target effect ^64^. Two complementary oligonucleotides were annealed and cloned into BbsI site of pSpCas9(BB)-2A-Puro (PX459) V2.0 (Addgene plasmid #62988) for co-expression with Cas9 using 5U of T4 DNA ligase, T4 DNA ligase buffer (1X) (Roche). 293T-LTR-GFP cells were transfected with the mix of equimolar ratio of PX459-sgRNA1 and PX459-sgRNA2 (Table S4). At 24h post-transfection, the medium was changed and 1μg/ml puromycin was added for selection and cells were cloned by serial dilution method.

## ACCESSION NUMBERS

The RNA-Seq data have been deposited on NCBI GEO under the accession number GSE123752.

---

**Figure S1:**
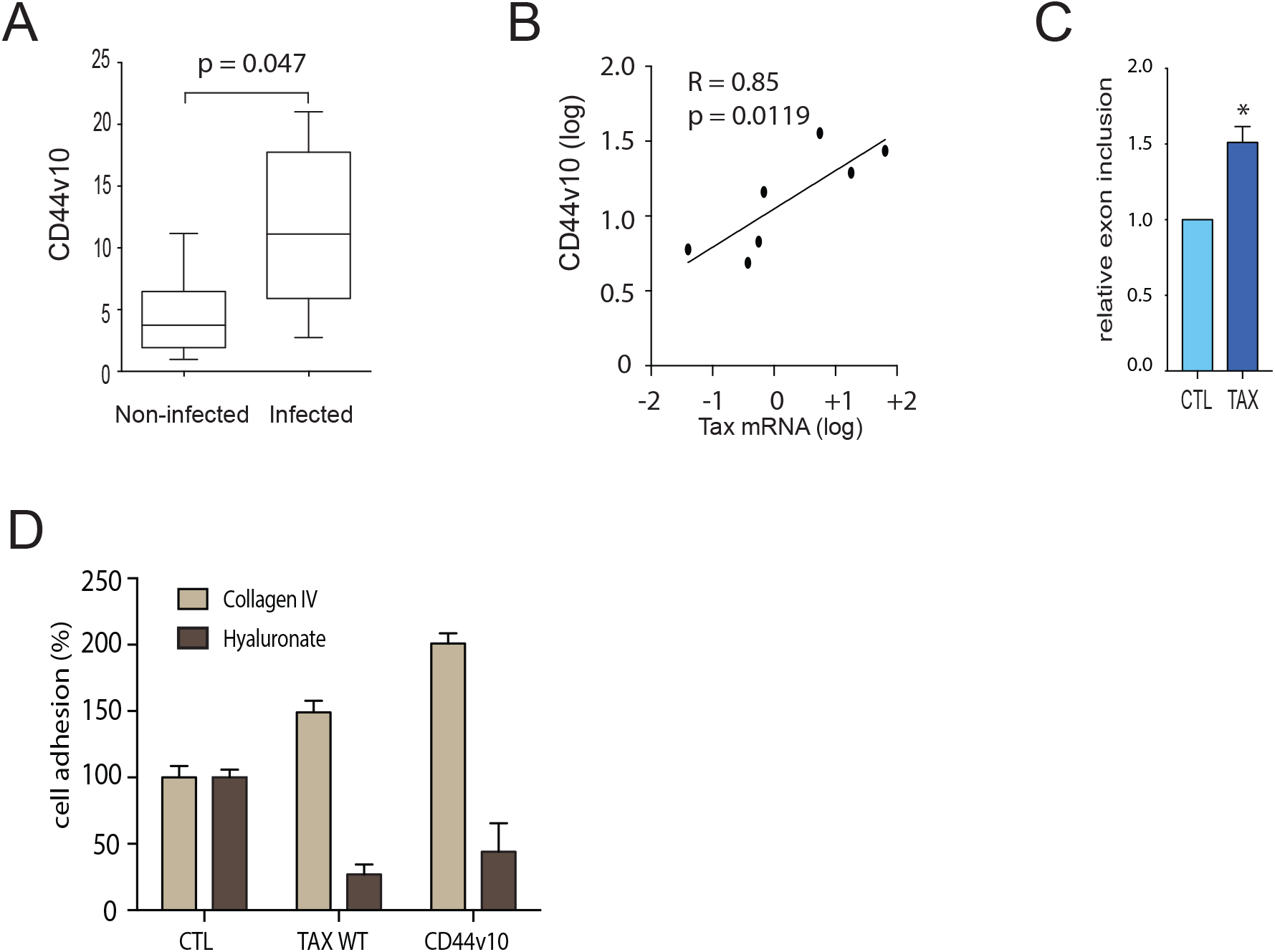
TAX-induced alternative splicing of CD44 in vivo and in vitro. (A) Expression of the splicing variant CD44v10 in HTLV-1 positive and negative cellular clones T-CD4+ derived from HAM/TSP patients (7 cellular clones in each category). CD44v10 mRNA were quantified by qRT-PCR from total RNA extrac_-2_ts. Median ±SD for non-infected vs infected clones were 11.1±6.87 vs 3.7±3.57; Mann Whitney test, p=0.047. (B) Positive correlation between TAX and CD44v10 mRNA expression in 7 infected CD4+ T-cell clones (Pearson correlation test). (C) qRT-PCR analysis of exon inclusion rate of CD44 exon v10 in 293 T-cells transiently transfected with pSG5M-TAX, pSG5M-M22 and empty pSG5M constructs. The relative exon inclusion of v10 corresponds to normalized v10 versus exon E16 (constitutive exon) expression levels. (D) Cell adhesion properties of cells transiently transfected with control vector, pSG5M-TAX and pCEP4-CD44V10 on plate surfaces coated with Hyaluronic acid and type IV Collagen. Histograms represent means ± s.d. of three independent experiments.

**Figure S2:**
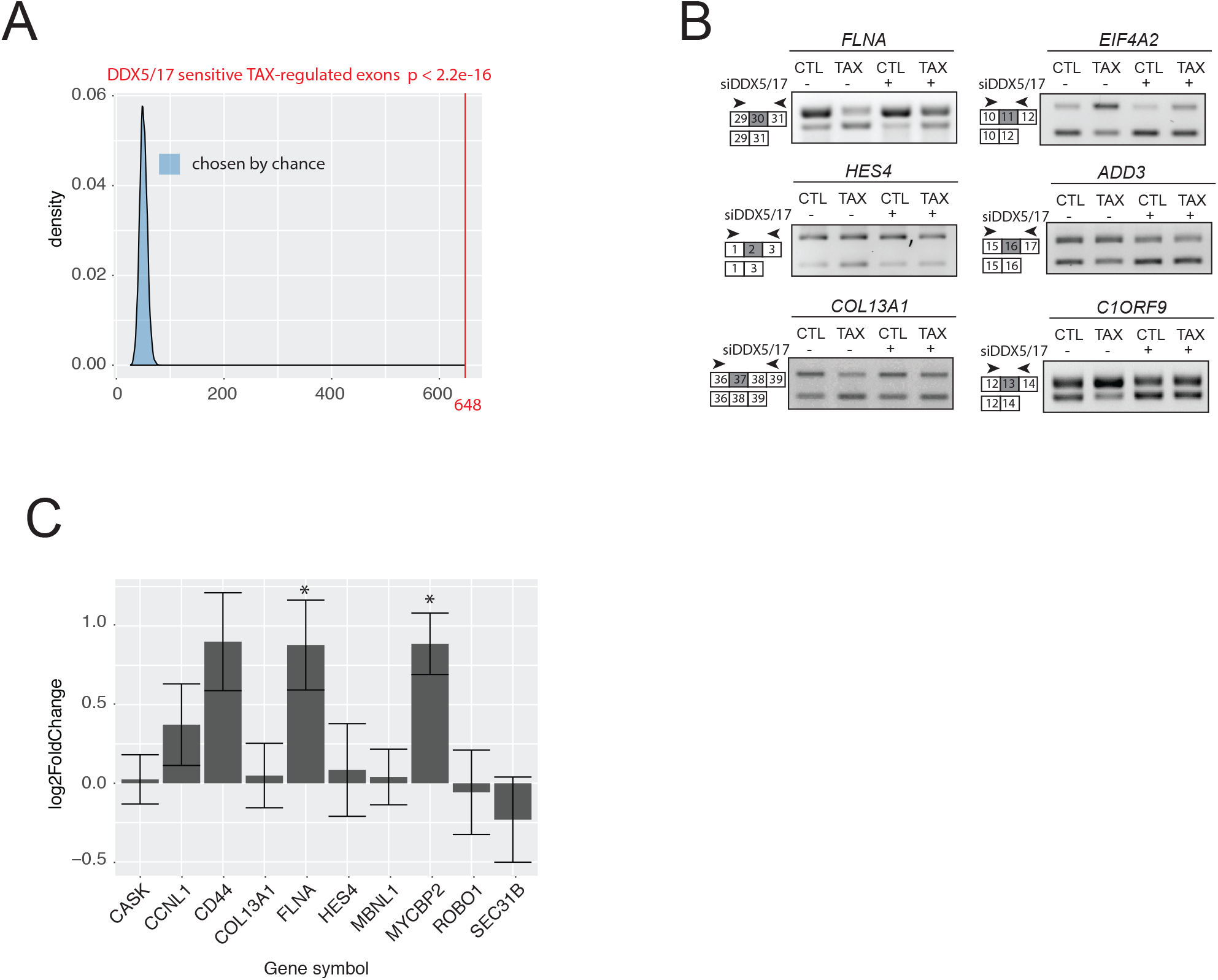
DDX5/17 expression regulate TAX splicing targets. (A) Bootstrapped distribution of DDX5/17 sensitive TAX-regulated exons and 648 randomly chosen exons (10000 repetitions) among overall expressed exons (blue). One Sample t-test p = 2.2e-16. (B) Validation of RNA-seq data using exon specific RT-PCR. (C) Fold change in gene expression of TAX-regulated exons responsive to DDX5/17 upon TAX expression. Values were obtained from DESeq2 analysis of RNA-seq data and expressed as Log2(FoldChange). Histograms represent means ± s.d. of three independent experiments. (*) p-val<0.05 (Mann Whitney test).

**Figure S3:**
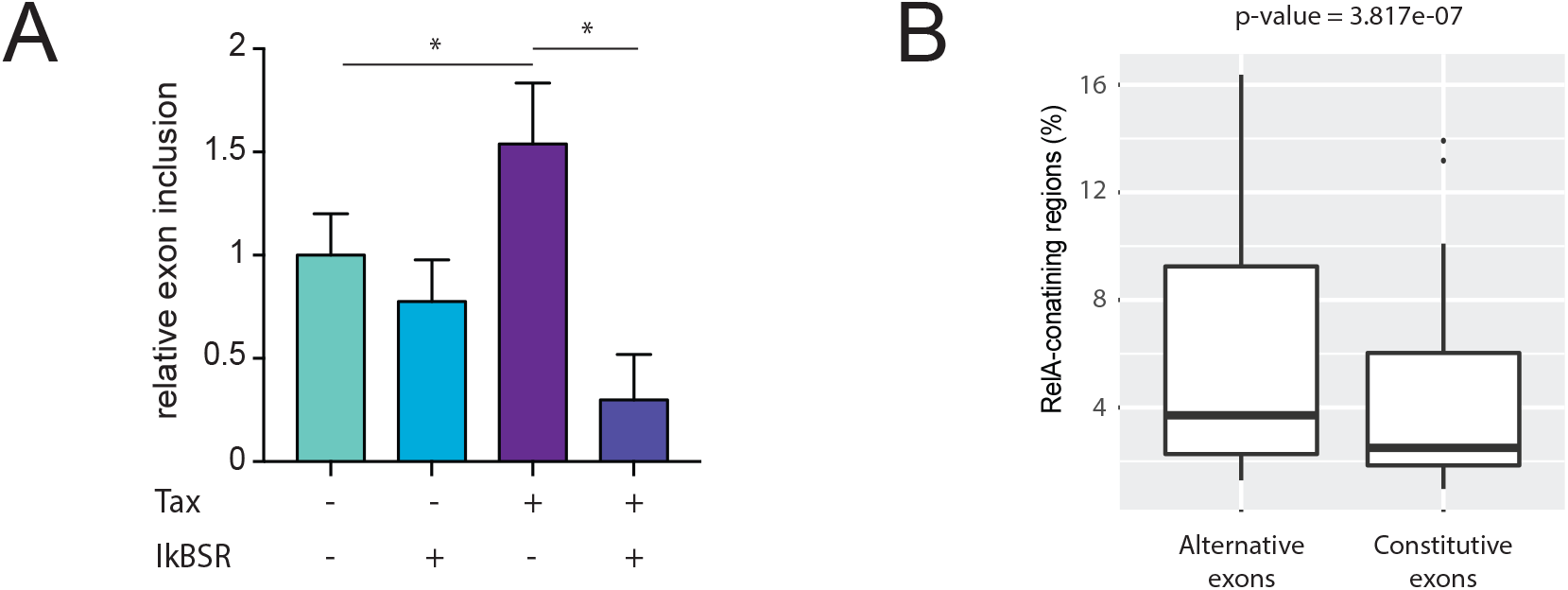
Chromatin tethering of RELA and DDX17 regulates alternative splicing. (A) TAX-induced exon inclusion of CD44 exon v10 relies on NF-kB activation. 293T-LTR-GFP cells were transiently transfected with the TAX vector along with the IkBSR expression vector and the corresponding empty control vectors. The inclusion rate of exon v10 was quantified by qRT-PCR. Histograms represent means ± s.d. of three independent experiments. (*) p<0.05 (Mann Whitney test). (B) Distribution of constitutive and alternative exons in RELA-enriched intragenic regions (3 kb). ChIP-seq datasets were analysed as detailed in method section. We excluded RELA peaks localized in intergenic regions and exons linked to specific events like pomoters, alternative first/last and mutually exclusive exons. The groups “Constitutive exons” and “Alternative exons” contained 41873 and 103000 exons, respectively. The window was fixed to 3 kb upstream and downstream of each exon coordinates. P-value was calculated using the Mann-Whitney test.

**Figure S4:**
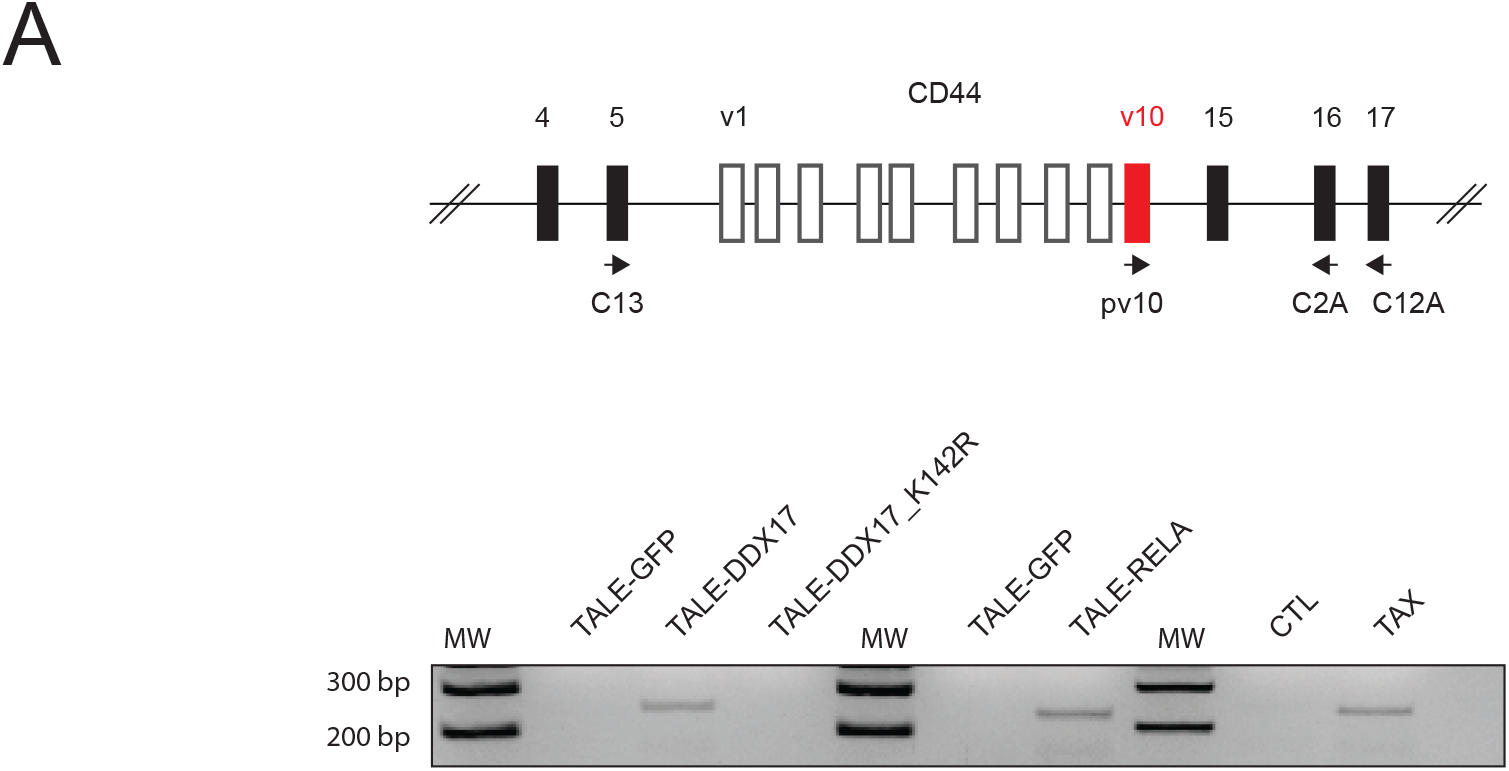
Nested RT-PCR analysis of CD44 transcripts expressed in TALE assays and in cells expressing or not TAX. Oligonucleotides were previously described (36). The first round of amplification consisted in 15 cycles of PCR with the primers C13 and C12A, the second round consisted in 35 cycles with primers pv10 and C2A. Final PCR products were resolved on Agarose gel (1%).

